# Evidence of autonomous neural specification for both brain and ventral nerve cord tissue in Annelida

**DOI:** 10.64898/2026.01.06.697942

**Authors:** Nicole B. Webster, Johnny A. Davila-Sandoval, Allan M. Carillo-Baltodano, Skyler Duda, B. Duygu Özpolat, Néva P. Meyer

## Abstract

Evolution of nervous systems is a long debated topic, and similar mechanisms of conditional neural specification linked to dorsal-ventral (D-V) axis formation across some taxa have been used to support homology. We tested for autonomous versus conditional neural specification in two distantly related annelids, *Capitella teleta* and *Platynereis dumerilii*, using blastomere isolations. Our results support previous work in *C. teleta* and further demonstrate that the autonomous specification of anterior neural tissue and for the first time in trunk neural tissue for both annelids. In *C. teleta*, we found evidence for conditional pro-neural and anti-neural signals for the VNC. Animal caps lacking vegetal macromeres at the 16-cell stage form a brain and a D-V axis but not a VNC while the addition of any single macromere rescues VNC fate. This suggests that animal micromeres other than 2d produce an anti-neural signal while a pro-neural signal is produced vegetally and that VNC specification is decoupled from D-V axis formation. Taken together, our study suggests possible conservation of autonomous specification of the brain and VNC within Annelida, raising interesting questions of how mechanisms controlling neural specification evolved in Spiralia.

## Introduction

The nervous system is a key innovation of animals whose origins and intricacies are still debated. Across Metazoa, there exist many different neural subtypes in varied circuit arrangements that enable the full breadth of behaviors and physiological functions. The large diversity of nervous system arrangements and functions offer intriguing opportunities to compare the molecular and cellular mechanisms controlling their formation across animals (Martín-Durán and Hejnol, 2021). Most studies of the nervous system have sensibly focused on a few key lineages, but with the advent of modern molecular tools, it has become feasible to widen our questions and investigations into less-well-known systems (Martín-Durán, 2023). It is only by expanding into ‘non-model systems’ that we can gain a fuller understanding of the diversity, development, and evolution of nervous systems.

Nervous system development is best understood in deuterostomes (e.g., vertebrates and hemichordates) and ecdysozoans (e.g., nematodes and arthropods). In vertebrates and flies, extrinsic factors involved in dorsal-ventral (D-V) axis formation, including Bone Morphogenetic Protein (BMP) signaling and activation of the MAPK cascade, are responsible for the correct development of the nervous system (Holley et al., 1995; Kuroda et al., 2004; Mieko Mizutani and Bier, 2008; Wilson and Edlund, 2001). This has led to an assumption that these two processes are intrinsically linked. Less is known about how nervous systems are specified in the third major bilaterian clade, Spiralia (∼Lophotrochozoa, e.g., annelids and mollusks) (Dunn et al., 2014; Martín-Durán and Hejnol, 2021).

There are striking differences in how a nervous system develops in spiralians. Recent results suggest that neural specification in Spiralia, at least in the episphere (larval ‘head’), may not require extrinsic factors. Instead, blastomere isolation data from the sedentary annelid *Capitella teleta* Blake et al., 2009 suggests that anterior neural specification, i.e. brain, is autonomous, occurring in individual cells without external signals such as morphogens (Carrillo-Baltodano and Meyer, 2017). Autonomous neural specification has only been reported in two other groups to date: the nerve cord but not brain of the ascidian *Halocynthia roretzi* (Minokawa et al., 2001) and the nerve net of ctenophores (Byrum et al., 2012). This is a striking difference in how the nervous system is specified developmentally in other groups and raises more general questions about the role of different developmental processes in cell fate specification.

Spiralians are an excellent group with which to study early stages of development. They have a stereotypical form of embryonic cleavage called spiral cleavage, thought to be ancestral for the group (Hejnol, 2010). The invariant nature of spiral cleavage allows comparative analyses of fates, body plans, and molecular mechanisms controlling development (Freeman and Lundelius, 1992; Henry, 2002). Individual blastomeres can be identified and tracked; at the 4-cell stage, the cells are named A, B, C, and D, and each gives rise to a different embryonic quadrant. These four macromeres divide along the animal-vegetal axis to generate the first tier of smaller cells on the animal side: the first-quartet micromeres (1a, 1b, 1c, 1d) or collectively 1q, while the cells on the vegetal side are first-quartet macromeres (1A, 1B, 1C, 1D) or 1Q. The cells continue dividing to give rise to subsequent tiers of micromeres, e.g., 2q, 3q, and 4q, until the onset of gastrulation (Conklin, 1897; Wilson, 1892). The stereotypical nature of spiral cleavage allows us to isolate specific cells or groups of cells to determine their role in downstream development and identify which cells are necessary or sufficient to produce specific structures.

In *C. teleta* and *Platynereis dumerilii*, fate-mapping has shown that the central nervous system (CNS) is formed by a few specific cells (Fig. 1; Ackermann et al., 2005; Meyer et al., 2010)). As in all other annelids and mollusks examined to date, the anterior ectoderm including the brain is formed from the first-quartet micromeres (1q; Ackermann et al., 2005; Clement, 1967; Damen and Dictus, 1996; Hejnol et al., 2007; Meyer et al., 2010; Render, 1991; Vopalensky et al., 2019; Weisblat et al., 1984). In annelids, the somatoblast responsible for forming the trunk neuroectoderm is derived from 2d or its equivalent (Ackermann et al., 2005; Matveicheva et al., 2024; Meyer et al., 2010; Shimizu and Nakamoto, 2015; Weisblat et al., 1984), while the source of the molluscan trunk neural tissue is less clear, but is generally some combination of 2q micromeres. The organizer is generally thought as a key piece in early developmental signaling to define affect neural tissue formation. In Spiralia, the organizer and is generally found in the D-quadrant. In *C. teleta*, blastomere deletion experiments demonstrated the organizing role of the micromere 2d for the correct formation of the D-V axis in the episphere, as well as formation of trunk mesoderm (Amiel et al., 2013). Whether the organizing nature of the D-quadrant arises from inherited determinants or is extrinsically specified varies between species (Freeman and Lundelius, 1992; Henry, 2014; Lambert, 2008; Seaver, 2017; Seudre et al., 2022), as does the specific D-quadrant organizer cell. In some mollusks and the annelid *Owenia fusiformis*, macromere 3D and its daughter micromere 4d function as the organizer (Goulding and Lambert, 2016; Henry and Perry, 2008; Henry et al., 2017; Seudre et al., 2022), while in annelids, micromere 2d (*C. teleta*, Amiel et al., 2013), sometimes in combination with 4d (*Tubifex tubifex*, Nakamoto et al., 2011), can also have organizing activity.

**Figure 1.**
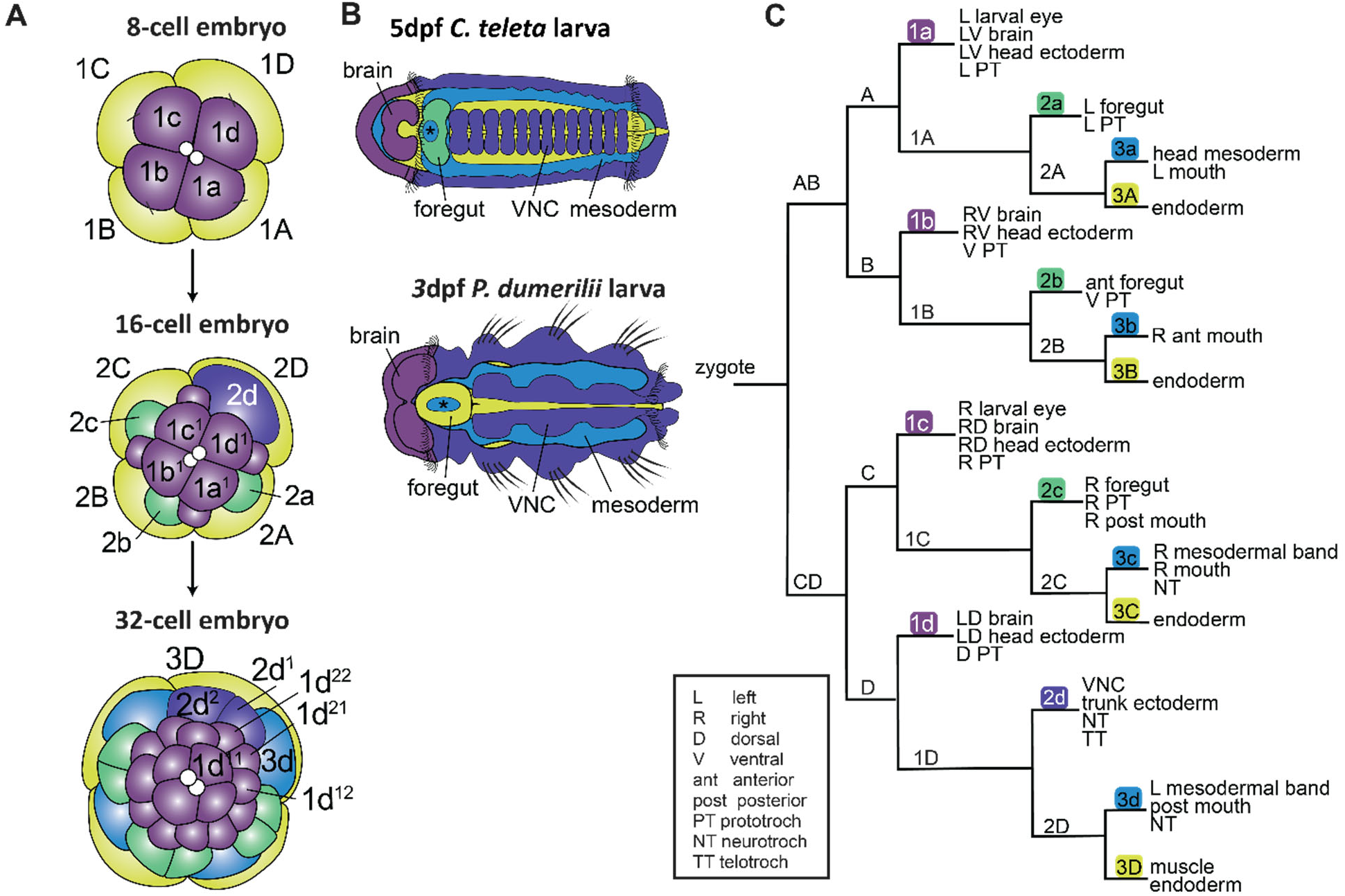
Fate map of specific structures. A) Micromere identity. B) Larval diagrams of *C. teleta* and *P. dumerlii.* B) Selective fate map of *C. teleta* modified from (Meyer et al., 2010). Colors correspond to cell identity. Violet: 1q, green: 2a, 2b, 2c; purple: 2d; yellow: 3Q; blue: 3q.

Blastomere isolations at the 8-cell and 16-cell stages in *C. teleta* previously suggested that the brain is specified autonomously while the VNC is specified conditionally (Carrillo-Baltodano and Meyer, 2017). Partial larvae derived from only 1q micromeres formed a “swimming head” with neural tissue including the neural marker *Ct-elav1* and FMRF-like but not serotonin (5HT) immunoreactivity. This suggests that 1q cells inherit maternal determinants that allow for rapid specification of the brain. In contrast, isolation of the -2Q animal cap at the 16-cell stage (1q^1^, 1q^2^, and 2q micromeres), which included 2d, resulted in partial larvae that did not express *Ct-elav1* in the trunk, suggesting conditional specification of the VNC. When -2Q animal caps were isolated at the 16-cell stage along with macromere 2D, which makes ectoderm, mesoderm and endoderm, *Ct-elav1* expression in the trunk was restored, suggesting that this macromere was able to provide extrinsic signals sufficient for neural specification of the VNC.

In this study we further explored the early role of specific blastomeres in neural specification and D-V axis formation in two annelids, *C. teleta* and *Platynereis dumerilii* (Audouin and Edwards, 1833). We also expanded on the types of isolations and the markers used to analyze tissue and axis formation. Most strikingly, we found that 2d alone *is* sufficient to produce trunk neural tissue in both annelids. This combined with our finding that isolated 1q micromeres form neural tissue in *P. dumerilii* suggests that the brain and VNC can be autonomously specified in both *C. teleta* and *P. dumerilii.* More broadly this further supports the hypothesis that specification of neural tissue and the D-V axis are decoupled.

## Results

### Neural specification

#### Anterior neural specification in *C. teleta* is autonomous in individual 1q micromeres

We previously demonstrated that 1q micromeres were capable of autonomous neural tissue formation when isolated together (1q: 1a+1b+1c+1d), but whether this applied to all quadrants individually was unclear. Here we tested if individual 1q micromeres (1a, 1b, 1c, or 1d) can produce neural tissue in isolation. Partial larvae derived from isolated 1a, 1b, 1c, or 1d micromeres were all very similar; they showed an anterior-posterior (A-P) axis with a posterior presumptive prototroch and no trunk or pygidium. This matches the ‘swimming head’ phenotype of 1q isolates but with less tissue (). Each quadrant produced a partial larva of a comparable size with multiple neural markers, including a single *Ct-elav1^+^* patch (1a: N = 4/4; 1b: N = 4/4; 1c; N = 7/7; 1d: N = 11/11), anti-acetylated Tubulin (ac-Tub; 1a: N = 5/5; 1b: N = 7/7; 1c; N = 5/5; 1d: N = 3/4), and FMRF-like immunoreactivity (FMRF-LIR; 1a: N = 5/5; 1b: N = 6/7; 1c; N = 3/5; 1d: N = 2/4). This further supports the hypothesis that brain tissue is autonomously specified in *C. teleta*.

#### VNC specification in *C. teleta* is autonomous in 1D-2D (2d) isolations

The entirety of the VNC as well as the ectoderm in the trunk and pygidium and the telotroch and neurotroch ciliary bands arise from the descendants of micromere 2d (Meyer and Seaver, 2010). We previously found that direct isolations of 2d are not feasible in *C. teleta*, likely because this is a very large cell that contacts all other cells at the 16-cell stage (Amiel et al., 2013). However, we were able to isolate 2d indirectly, by first isolating its parent cell (1D), then removing its larger, yolk-filled sibling cell 2D after division. These 1D-2D partial larvae were generally elongate with an A-P axis; they had no episphere features but there was a pronounced pygidium on one end (Fig. 3). Cilia were generally present on one end of the elongated partial larvae (N = 19/24), likely the pygidium, but the cilia were not clearly arranged into a ciliary band, i.e., the telotroch (Fig. 3H”-J”). Sometimes patches of cilia were also present in the ‘trunk’ (N = 5/24), which may represent a partial neurotroch. A lack of phalloidin staining suggests an absence of muscles, and likely mesoderm (N = 16/16), as expected. Both FMRF-LIR (N = 7/7) and *Ct-elav1* (N = 23/26) were present in the 1D-2D partial larvae, showing the presence of neural tissue. Clusters of cells expressing *Ct-elav1* were generally seen throughout the trunk but they were not confined to any one side of the animal, i.e., the ventral side. These results suggest that 2d is capable of autonomously producing neural tissue in the absence of other blastomeres including those that generate mesoderm.

**Figure 2.**
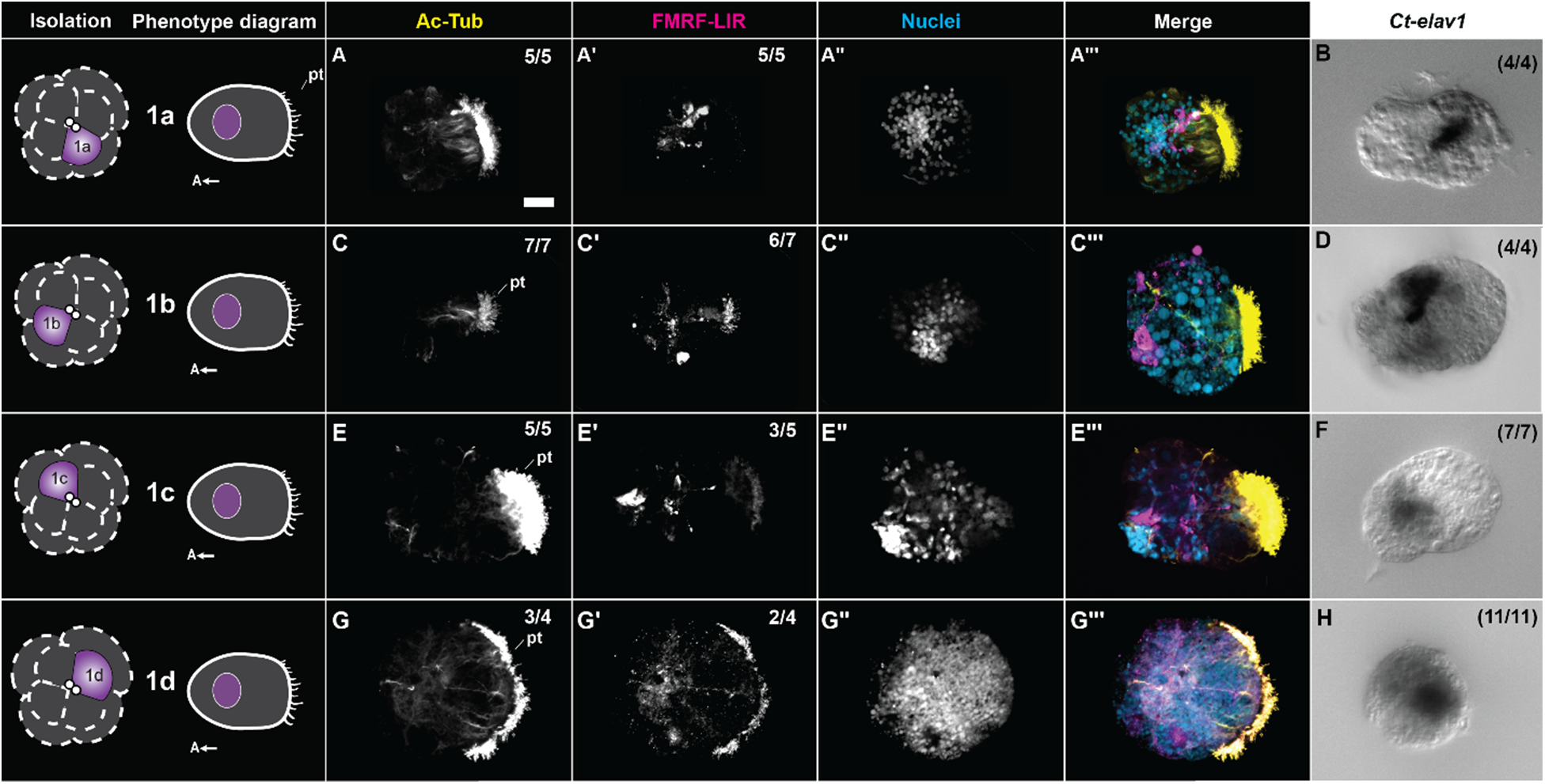
Phenotypes of 1a, 1b, 1c, and 1d isolates from 8 cell embryos in *C. teleta*. A,B) 1a isolates. C,D) 1b isolates. E,F) 1c isolates. G,H) 1d isolates. A,C,E,G) Acetylated tubulin immunoreactivity (Ac-Tub; yellow in merged images). A‵,C‵,E‵,G‵) FMRF-LIR (magenta in merged images). A‵‵,C‵‵,E‵‵,G‵‵) Hoechst nuclei (cyan in merged images). A‵‵‵,C‵‵‵,E‵‵‵,G‵‵‵) Merged fluorescent image. B,D,F,H) *Ct-elav1* ISH. Images with the same letter are the same individual. Left-most column: Isolation and phenotype diagrams. Top right numbers in figure panel: number of isolates with shown neural phenotype out of total. AcT: ac-tub neural tissue; pt: prototroch; scale bar: 20 µm.

**Figure 3.**
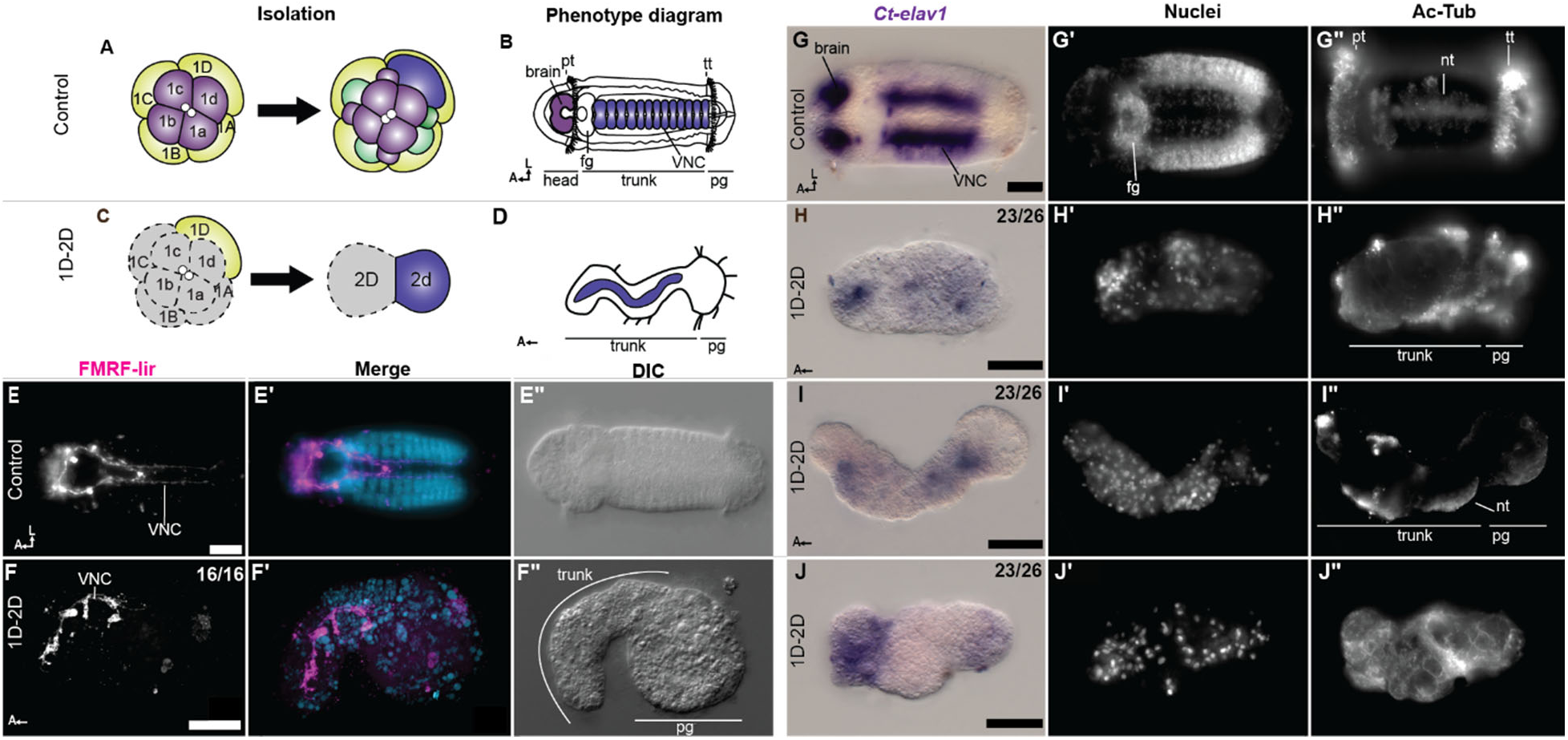
Phenotypes of control and 1D-2D (2d) isolates from 16 cell embryos in *C. teleta*. A) Control isolation diagram. B) Control CNS diagram, colours represent the source cells in A. C) 1D-2D isolation diagram. D) 1D-2D isolation phenotype diagram. G, H, I, J) *Ct-elav1* ISH. G) Control animal. H-J) representative 1D-2D partial larvae. G‵-J‵) nuclei. G‵‵-J‵‵) Ac-Tub. E-F) FMRF-LIR. E) Control animal. F) 1D-2D partial larva. E,F) FMRF-LIR. E‵,F‵) FMRF-LIR (magenta) and Hoechst nuclear (cyan) merged images. E”, F”) DIC images. Images with the same letter are the same individual. nt: neurotroch; pg: pygidium; pt: prototroch; tt: telotroch; scale bars: 50 µm.

#### Any 2Q macromere can rescue the VNC in -2Q animal caps

We previously showed that isolation of animal caps including 2d at the 16-cell stage (-2Q animals) resulted in an elongated partial larvae with an anterior brain but no VNC. Addition of the 2D macromere, which generates mesoderm and endoderm on the left side of the trunk, rescued VNC fate in the trunk (Carrillo-Baltodano and Meyer, 2017). To better understand how cells from different quadrants affect neural specification in the trunk, we isolated -2Q animal caps with either macromere 2A, 2B, or 2C. In *C. teleta*, descendants of macromeres 2A and 2C generate endoderm plus anterior mesoderm or right trunk mesoderm, respectively, while macromere 2B generates endoderm and not much if any mesoderm. Furthermore, blastomere 2d is thought to act as the D-V organizer for at least the episphere (Amiel et al., 2013). The resulting partial larvae were similar in phenotype to -2Q animal caps + 2D; all showed a clear A-P axis and the vast majority had a D-V axis with a head, brain, foregut, stomodeum, prototroch, and telotroch (Fig. 4). Each macromere was sufficient to restore the VNC (*Ct-elav1*; -2Q+2A (N = 17/22), - 2Q+2B (N = 16/22), -2Q+2C (N = 23/28), or -2Q+2D (N = 16/16), but there were variations in the degree of ‘rescue’. Assessing the amount of *Ct-elav1* and the presence of a ventral midline (determined by the presence of a gap between two lateral bands of *Ct-elav1* expression) suggests that -2Q+2A had the weakest VNC rescue: the majority of these partial larvae had only weak, short (∼1 ganglion) regions of *Ct-elav1* (N = 6/8), and none showed a midline. The -2Q+2C had the next weakest rescue where only 60% showed evidence of a midline (N = 4/7) and the majority of had less than 5 ganglia as determined by *Ct-elav1* expression (N = 5/8). The -2Q+2B partial larvae were 75% likely to show a midline (N = 10/13) and the majority showed at least 5 ganglia with *Ct-elav1* (N = 7/10). In contrast, all - 2Q+2D animals had *Ct-elav1*, ∼90% of - 2Q+2D show a midline (N = 16/18) and a long *Ct-elav1^+^* expression band (N = 16/18 with more than 5 ganglia). While these details are not sufficient to tease apart the signaling differences between the 2Q macromeres, all four vegetal macromeres at the 16-cell stage or their descendants can produce proneural signals and these signals are not localized to cells involved in D-V axis formation (i.e., the D quadrant) or to cells that generate mesoderm (e.g., only 2A, 2C, and 2D).

**Figure 4.**
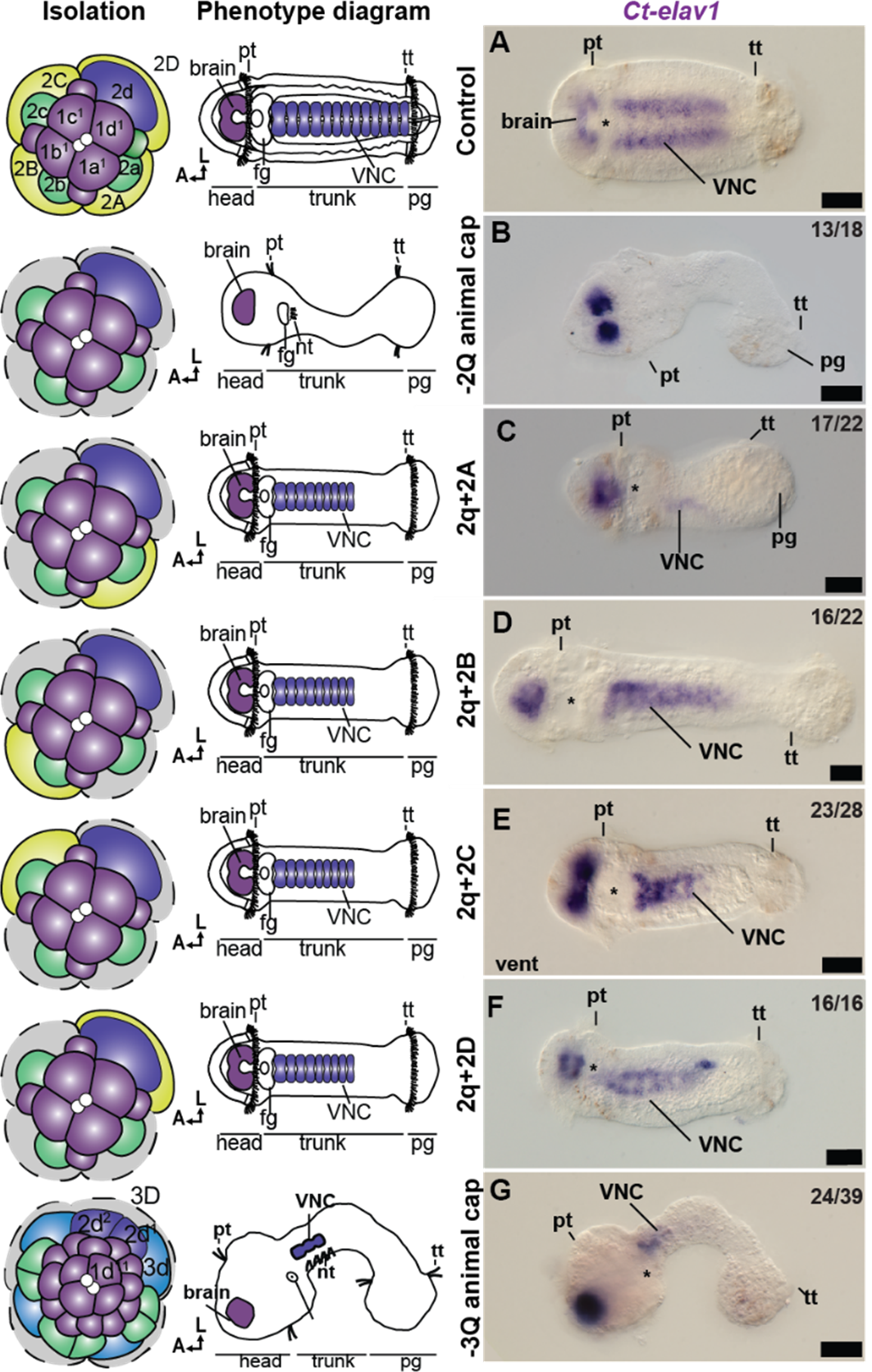
*Ct-elav1* expression in partial *C. teleta* embryos including the 2q animal cap (ventral view). A) Representative control embryo. B) -2Q animal cap. C) -2Q animal cap including 2A. D) -2Q animal cap including 2B. E) -2Q animal cap including 2C. F) -2Q animal cap including 2D G) -3Q animal cap. Left-most column: diagram of isolation. Top right numbers in figure panel: number of isolates with shown phenotype out of total. Asterisk: position of the mouth opening. A: anterior; D: dorsal; fg: foregut; nt: neurotroch; pg: pygidium; pt: prototroch; tt: telotroch; VNC: ventral nerve cord. Scale bars: 50 µm.

#### -3Q animal caps have a reduced VNC

To test if partial larvae isolated after birth of the 3q micromeres could produce VNC, we isolated the animal cap at the 32-cell stage (28-cell animal cap consisting of 1q+2q descendants and 3q micromeres; - 3Q animal cap). -3Q animal cap partial larvae looked very similar to the animal caps isolated at the 16-cell stage with one macromere, except they had a narrower trunk and less VNC tissue (Fig. 4G). They had a clear A-P axis with an episphere and pygidium, and most had a clear D-V axis with a neurotroch (N = 10/14) and a ventral midline, although only some showed evidence of trunk mesodermal bands (nuclear localization; N = 5/14). They showed both episphere (N = 39/39) and trunk *Ct-elav1* expression (Fig. 3G *Ct-elav1*; N = 24/39), albeit only in less than 5 anterior ganglia (N = 17/20). The presence of trunk neural tissue in partial larvae generated from -3Q animal caps suggests that conditional signals at the 16-cell stage, both proneural signals from the macromeres and anti-neural signals from the micromeres, have stopped by the 32-cell stage.

### Anatomy of -2Q animal caps in *C. teleta*

While the partial larvae derived from -2Q animal caps isolated at the 16-cell stage do not express neural markers in the trunk, they do elongate and seem to form an A-P axis with an episphere, trunk, and pygidium. We previously reported a lack of a D-V axis, foregut, and mouth (Carrillo-Baltodano and Meyer, 2017). However, based on recent findings that the 2d blastomere is likely the D-V axial organizer in *C. teleta* (Lanza and Seaver, 2020, 2018), we assessed the expression of additional D-V and tissue-specific markers in partial larvae generated by -2Q animal caps (Fig. 5).

**Figure 5.**
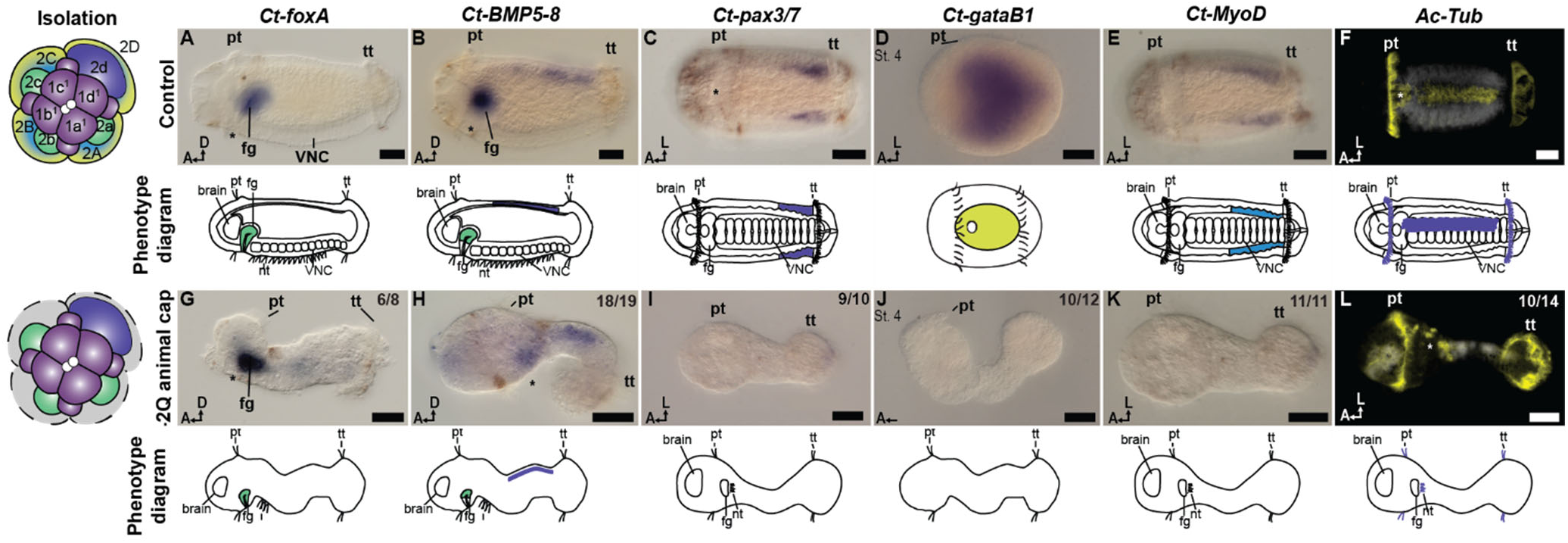
Gene expression in partial *C. teleta* larvae including the -2Q animal cap. A-F) Control embryos. G-L) -2Q animal cap. A,G) *Ct-foxA* ISH at stage 6. B,H) *Ct-BMP5-8* ISH at stage 6. C,I) *Ct-pax3/7* ISH at stage 6. D,J) *Ct-gataB1* ISH at stage 4. E,K) *Ct-MyoD* ISH at stage 6. F,L) Ac-tub (yellow) and Hoechst (white) at stage 6. Left-most column: diagram of isolation. Top right numbers in figure panel: number of isolates with shown phenotype out of total. Line drawings: diagram of above phenotype. Asterisk: position of the mouth opening. A: anterior; D: dorsal; L; left; fg: foregut; pt: prototroch; tt: telotroch. Scale bars: 50 µm.

#### -2Q animal caps have a D-V axis

*Ct-foxA* is normally expressed in the ventrally-positioned foregut at stage 6, which is located immediately dorsal to the stomodeum (Fig. 5A, Boyle and Seaver, 2008). The ectodermal foregut is formed by cells from the 2a, 2b, and 2c lineages (Meyer et al., 2010). In -2Q animal cap partial larvae, *Ct-foxA* was expressed in the anterior region of the elongated trunk, posterior to the prototroch and on the same side as the mouth opening (N = 6/8), suggesting a D-V axis is formed in the absence of 2Q macromeres (Fig. 5G).

*Ct-bmp5-8* is normally expressed on the dorsal side of the foregut as well as in two dorsolateral stripes along the trunk ectoderm (Fig. 5B, Webster et al., 2021). In the partial larvae derived from -2Q animal caps, *Ct-bmp5-8* is expressed in both a ventral and dorsal ectodermal domain. The first is anterior, just posterior to the prototroch, and on the same side as the mouth opening (N = 18/19; Fig. 5H); the second domain is opposite and more posterior, suggesting that this is the dorsal domain, although there is no sign of two separate dorsal stripes (N = 14/19). This expression pattern further supports D-V axis formation in partial larvae generated by -2Q animal caps.

We initially examined *Ct-pax3/7* expression as a D-V marker in the partial larvae as this gene is normally expressed in two ventrolateral ectodermal bands that reduce to the region near the posterior growth zone at later larval stages (Fig. 5C and Seaver et al., 2012). In partial larvae derived from -2Q animal caps, no *Ct-pax3/7* expression was observed (N = 10/10; Fig. 5I). This could indicate a lack of ventrolateral ectoderm. However, Pax3/7 homologs are thought to be involved in later stages of neurogenesis in at least some spiralians (Navet et al., 2017), and the lack of *Ct-pax3/7* in the partial larvae likely results from a lack of neural specification leading to an absence of neurogenic markers.

We also used ac-Tub immunolabeling to look for ciliated cells that might represent remnants of the neurotroch, a ciliary band that normally runs along the ventral midline in the trunk (Fig. 5F). We found that -2Q animal cap partial larvae had patches of external cilia in the trunk that were on the same side as the stomodeal opening (N = 10/14; Fig. 5L), supporting the presence of a neurotroch and ventral, non-neural ectoderm. We do not think that these cilia are the external cilia that surround the mouth opening, and are instead true neurotrochal cilia, as the cilia near the mouth in wild-type larvae come from third-quartet micromeres (Meyer et al., 2010), which were removed in our experiment. A re-examination of the partial larvae derived from -2Q animal caps in Carrillo-Baltodano and Meyer (2017), showed that most of these larvae also had a mouth opening, although a foregut lumen was not detectable despite the presence of foregut markers (N = 42/52). Taken together, these molecular and morphological markers indicate that there is an ectodermal D-V axis but no VNC in the trunk of larvae derived from -2Q animal caps in *C. teleta*.

#### -2Q animal caps lack endoderm and mesoderm

At the 16-cell stage in *C. teleta*, mesoderm and endoderm are largely generated by progeny of the macromeres (Meyer et al., 2010), which are the cells being removed in the -2Q animal caps. In order to confirm that these partial larvae lack mesodermal and endodermal tissue, we examined two markers, *Ct-myoD* and *Ct-gataB1* (Boyle and Seaver, 2008). *Ct-gataB1* is most visible in the endoderm at stage 4 (Fig. 5D), and when control animals were at stage 4, there was no *Ct-gataB1* expression observed in -2Q animal cap partial larvae (N = 10/12; Fig. 5J), suggesting a lack of endodermal tissue in these animals. Interestingly, the stage 4 partial larvae were already elongated whereas control animals were still ovoid in shape, which may be due to the lack of the large yolky macromeres during gastrulation, possibly accelerating the process of convergent-extension. *Ct-myoD* is expressed in the left and right mesodermal bands and has a role in muscle development (Boyd and Seaver, 2023). In stage 6 controls, *Ct-myoD* was expressed in the posterior region of both mesodermal bands (Fig. 5E), but in -2Q animal caps, there was no clear *Ct-myoD* expression (N = 11/11; Fig. 5K) suggesting a lack of mesodermal tissue as expected.

### Blastomere isolations in *P. dumerilii* recapitulate the pattern of autonomous neural specification in *C. teleta*

To test whether the results in *C. teleta* (Sedentaria), can be generalized more broadly, we performed blastomere isolations in the distantly-related annelid *P. dumerilii* (Errantia; Fig. 6) (Struck et al., 2011; Weigert and Bleidorn, 2016). Errantia + Sedentaria comprise Pleistoannelida and members of these clades diverged no later than the Early Ordovician (∼490 Ma; Parry et al., 2014), with distinctly different lifestyles (errant vs sedentary) and associated morphologies (tubes vs swimming parapodia). Both *C. teleta* and *P. dumerilii* have unequal spiral cleavage with early blastomeres displaying very similar fates (Ackermann et al., 2005; Meyer et al., 2010). In *P. dumerilii*, isolated 1q micromeres developed into swimming heads, similar to what we previously described for *C. teleta* (Carrillo-Baltodano and Meyer, 2017). They all showed a clear A-P axis, with a posteriorly-localized prototroch and anteriorly-localized neural markers, but no eyes (N = 24/24; Fig. 6D–F), which contrasts with formation of eyes in some 1q partial larvae in *C. teleta*. Specifically, 1q partial larvae had neurites and cilia labeled with ac-Tub (N = 24/24), FMRF-LIR (N = 11/12), and *Pdu-elav1* expression (N = 2/2) but not 5HT (N = 0/12), similar to *C. teleta*. These data suggest that *P. dumerilii* can also produce anterior neural tissue autonomously.

**Figure 6.**
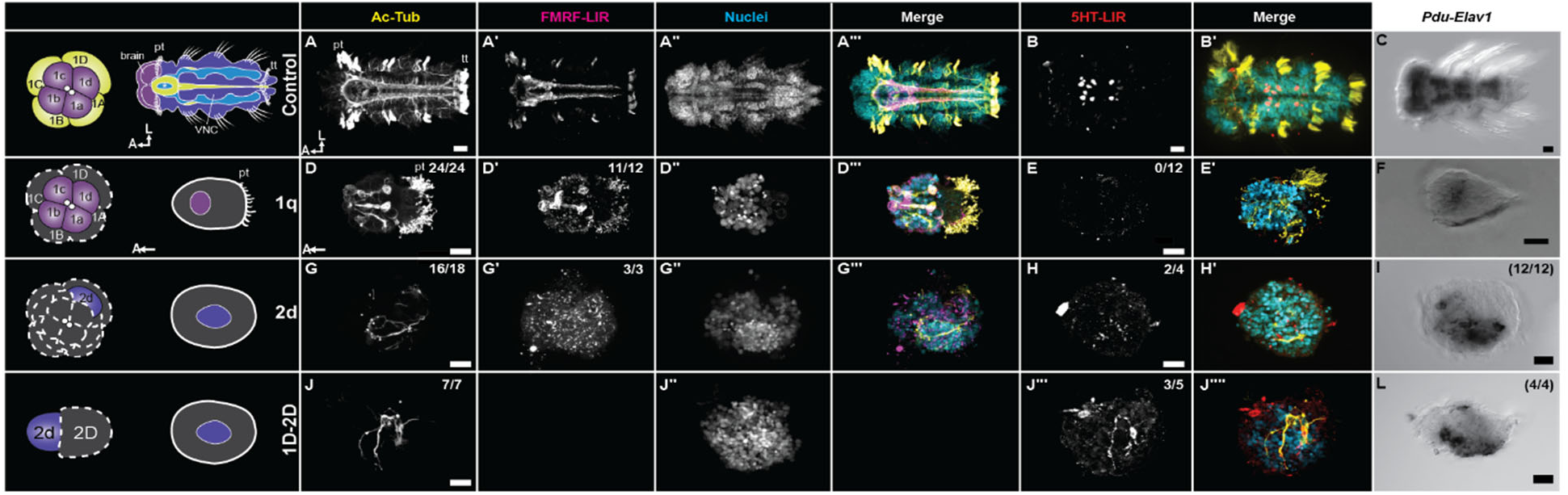
Phenotypes of *P. dumerilii* isolates. A,B,C) Control. D,E,F) 1q isolates. G,H,I) 2d direct isolates. J,K,L) 1D-2D isolates (2d indirect isolations). A,D,G,J) Acetylated tubulin immunoreactivity (yellow in merged images). A‵,D‵,G‵) FMRF-LIR (magenta in merged images)‵. A‵‵,D‵‵,G‵‵,J‵‵) Hoechst nuclear marker (cyan in merged images). A‵‵‵,D‵‵‵,G‵) Merged fluorescent image. B,E,H,J‵‵‵) Serotonin (5-HT) immunoreactivity (red in merged images) B‵,E‵H‵J‵‵‵‵) Merged fluorescent image. C,F,I,L) *Pdu-elav1* ISH. Images with the same letter are the same individual. Left-most column: diagram of isolation. Top right numbers in figure panel: number of isolates with shown neural phenotype out of total. A: anterior; D: dorsal; L; left; pt: prototroch; tt: telotroch; scale bars: 20 µm.

Unlike in *C. teleta* we were able to directly isolate the 2d micromere in *P. dumerilii*. However, the 2d partial larvae did not elongate as they do in *C. teleta*, and instead appeared more as an ovoid ball of cells with no external features: no chaetae, only disorganized cilia, and no distinguishable axes. However, 2d isolations did have neural tissue (Fig. 6G–I). This included neurites and cilia labeled with ac-Tub (N = 16/18), FMRF-LIR (N = 3/3), *Pdu-elav1* expression (N = 12/12), and unlike the episphere, some 5HT (N = 2/4). Although the neural tissue was sometimes confined to a single side of the embryo, this was not consistent. We also carried out the 1D-2D isolations for comparison and found that 1D-2D partial larvae were generally indistinguishable from 2d partial larvae with no morphologically-distinguishable axes or external features. These partial larvae also expressed the neural markers (Fig. 6J–L) ac-Tub (N = 7/7), *Pdu-elav1* (N = 4/4), and 5HT (N = 3/5). This supports using 1D-2D as a proxy for a 2d direct isolation in *C. teleta* as there were no phenotypic difference between the two isolations in *P. dumerilii*. It also suggests that the failure to elongate of 2d isolates was not due differences in isolation procedures between *C. teleta* and *P. dumerilii*. Taken together, these data suggest that 2d also can form VNC tissue autonomously in *P. dumerilii*.

## Discussion

### Autonomous and conditional neural specification in Pleistoannelida

Our primary result is the clear presence of neural tissue in partial larvae derived from just a single cell—micromere 2d—in two phylogenetically-distinct species of annelids *C. teleta* (Sedentaria) and *P. dumerilii* (Errantia). Furthermore, our data shows that isolated 1q micromeres make neural tissue in both species, supporting autonomous specification via inherited determinants for the brain and VNC in Pleistoannelida. While individual micromeres at the 8- and 16-cell stages can produce neural tissue in isolation, there is also evidence that conditional signaling during cleavages stages affects neural fates. For the brain, 1q isolates in *C. teleta* (Carrillo-Baltodano and Meyer, 2017) and *P. dumerilii* lacked 5HT, suggesting that some neural subtypes in the brain require extrinsic signaling. Furthermore, we showed that pro-neural and anti-neural signaling from other blastomeres are involved in VNC specification. Isolating micromere 2d with other animal micromeres (-2Q partial larvae) at the 16-cell stage blocks VNC but not brain formation in *C. teleta.* Moreover, any single vegetal 2Q macromere, not just macromere 2D, is sufficient to rescue VNC fate when isolated with the animal micromeres at the 16-cell stage. Signaling from other blastomeres (pro- and anti-neural) is thus necessary to maintain balance in neural fate specification in the trunk in complete larvae. Overall, we have demonstrated that both autonomous and conditional signaling during cleavage stages regulate both head and trunk neural specification in two species of annelids.

We did not identify the specific genetic mechanisms or signaling centers for neural specification; however, our data suggest a few hypotheses in *C. teleta*. Firstly micromeres 1q^2^ and/or 2a–c produce anti-neural signals for the VNC but not brain. -2Q animal caps consist of micromeres 1q^1^, 1q^2^, and 2q, and they generate a brain but not a VNC. Neural fate may be largely autonomously determined in 1q^1^ cells, making any anti-neural signals from 1q^2^ or 2q insufficient to alter their trajectory, explaining why anterior neural tissue was not affected. Interestingly, both 1q^2^ and 2a–2c micromeres primarily produce ectodermal tissues, the prototroch and foregut, respectively, although micromeres 2a–2c also generate interstitial cells in the trunk, which were previously interpreted as ectomesoderm (Meyer et al., 2010). This would suggest that the anti-neural signal comes from cells that generate non-neural ectoderm.

Our data also suggest that 2Q macromeres at the 16-cell stage produce a pro-neural, vegetally-localized signal for the VNC that is not restricted to the D-quadrant. By adding any single 2Q macromere to the - 2Q animal cap, the VNC could be rescued. The timing of this signal is restricted by the fact that isolating the -3Q animal cap (after the birth of the 3q micromeres), just 1 hour later, resulted in development of a small amount of VNC tissue. This could indicate that by the 32-cell stage, the anti-neural signal has decreased or the pro-neural signal has already had an effect. Alternatively, the pro-neural signal could be produced by the 3q micromeres after their birth. Either way, neural specification appears to occur well before gastrulation in *C. teleta*. Surprisingly, the pro-neural signal is not localized to the D quadrant macromere, 2D, as any of the four macromeres can rescue VNC fate. This suggests a decoupling of neural fate and the organizing activity of the D quadrant found in many spiralians (discussed below), at least in the trunk. Micromere 2d organizes the D-V axis in at least the episphere in *C. teleta* (Amiel et al., 2013). Finally, the pro-neural signal does not come from mesodermal precursors since macromere 2B, which does not generate mesodermal derivatives in *C. teleta* (Meyer et al., 2010), can rescue VNC fate. This is interesting since mesoderm plays an important role in inducing neural tissue in vertebrates and insects (Beddington, 1994; Jones and Woodland, 1989; Kiecker and Niehrs, 2001; Lynch and Roth, 2011).

Lastly, mechanisms of neural specification in the episphere and trunk are different. While both the brain and VNC can be autonomously specified, there appear to be additional pro- and anti-neural signals for the VNC while in the brain only neural subtypes (e.g., 5HT^+^ neurons) rely on conditional signals. This fits with the larger picture of larval evolution suggesting that there is a decoupling of episphere and trunk formation that may have facilitated larval and life cycle evolution (Carrillo-Baltodano and Meyer, 2017; Martín-Zamora et al., 2023). Indeed, even the definition and make up of a brain varies between groups (Nielsen, 2005; Sumner-Rooney and Sigwart, 2018). For example, the larval apical organ might be considered an ancestral eumetazoan brain, but is not homologous to the adult brain and is lost at metamorphosis (Moroz, 2009; Nielsen, 2005). If episphere and trunk neural tissue have different regulatory mechanisms, perhaps they should be treated differently when investigating the homology of CNSs across Bilateria (Martín-Durán et al., 2018; Nanglu et al., 2023).

Whether this neural specification pattern is conserved across spiralians remains to be seen, but we think that this is unlikely. Spiralia contains organisms with a wide array of nervous system arrangements, and existing data on the role of BMP on neural tissue formation in spiralians shows a lack of developmental conservation (Lyons et al., 2020; Tan et al., 2022; Webster et al., 2021). Recent evidence on the role of BMP signaling in different annelids points to the possible evolution of separate mechanisms depending on the type of spiral cleavage (equal vs unequal cleavage) (Carrillo-Baltodano et al., 2025; Seudre et al., 2022). Ultimately, identification of the autonomous and conditional signals for brain and VNC specification in different species will help us understand if these two systems evolved in parallel or if there was a common molecular mechanism controlling both that then.

### Separation of neural specification and D-V axis formation in Annelida

Elucidating the molecular and cellular mechanisms controlling neural specification in a wide array of animal clades is crucial to reconstructing the evolutionary trajectory of nervous systems and their relationship to the D-V axis. Although there has been recent progress in comparing the molecular control of D-V axis formation across spiralian taxa, the control of neural specification in spiralians is relatively unknown.

For comparison, in some vertebrate and arthropod taxa studied to date, gradients of opposing signaling systems, e.g., BMP signaling versus EGF/FGF signaling, organize fates relative to one another to form the D-V axis. One such fate is that of the neuroectoderm, which arises on one side of the D-V axis due to upregulation of the MAPK cascade and inhibition of BMP signaling, although other signals such as Wnt also play a role depending on the species and position along the A-P axis (Lynch and Roth, 2011; Stern, 2005). The molecular similarities in neural specification between vertebrates and insects invigorated the hypothesis that a CNS evolved once in the last common ancestor of bilaterians (Arendt et al., 2008; Arendt and Nübler-Jung, 1994). However, recent data on spiralians, including work from our lab, have found varied to no involvement of MAPK activation and inhibition of BMP signaling in neural specification, and the relationship to organization of fates along the D-V axis is unclear (Allan, 2024; Carrillo-Baltodano, 2019; Kuo and Weisblat, 2011; Lambert et al., 2016; Pfeifer et al., 2014). For example, we previously found that upregulating BMP signaling during cleavage stages in *C. teleta* may promote a D- and C- quadrant identity at the expense of a B-quadrant identity but does not repress formation of brain or VNC tissue. This would suggest that BMP signaling in *C. teleta* is not an anti-neural signal as in some other animals (Webster et al., 2021). Other data from *C. teleta* have shown that Activin signaling via SMAD 2/3 but not BMP signaling via SMAD1/5/8 organizes the D-V axis in the episphere and trunk but does not repress neural tissue formation. This D-V organizing signal is likely generated by micromere 2d as its ablation abrogates D-V axis formation in the episphere (Amiel et al., 2013; Lanza and Seaver, 2020, 2018). Overall, these data demonstrated that blocking Activin signaling or deleting 2d resulted in a loss of the D-V axis but not neural tissue. Carrillo-Baltodano *et al*. (2025) suggest that this is an annelid-specific molecular mechanism that can be explained by developmental systems drift. They show that BMP does not play a role in D-V axis signaling in annelids with unequal spiral cleavage (*C. teleta* and *P. dumerilii*) but is involved in D-V axis formation in species with equal spiral cleavage (*Owenia fusiformis* and *Spirobranchus lamarcki*). Our data here complement these findings; isolated -2Q animal caps express non-neural ectodermal gene markers in an ordered manner, *Ct-foxA* ventrally and *Ct-bmp5-8* dorsally, but not neural markers, suggesting formation of the D-V axis in the absence of the VNC. Taken together, this would suggest that in *C. teleta* D-V organizer signaling is not responsible for restricting neuroectoderm to one region of the D-V axis. Instead, a combination of inherited factors plus additional pro-neural and anti-neural signals, which may not even originate from one quadrant, establish the neuroectoderm. This raises an intriguing question of how the brain and VNC in *C. teleta* become localized to one region along the D-V axis. If these two developmental features are ancestrally decoupled in Spiralia it should be considered as a key piece of evidence to explain the origin(s) of a centralized nervous system.

### *P. dumerilii* and *C. teleta* partial embryos developed differently

While the neural specification patterns we tested were conserved between these two species, a number of concerns limited the interpretation of *P. dumerilii* blastomere isolations. As a result, data from -2Q animal caps and other late isolations were inconsistent and not included here. In particular, *P. dumerilii* partial larvae never had eyes or an elongated body shape. Without the elongated body shape, it was more difficult to control for embryos that did not gastrulate properly or to even determine an A-P axis. We believe this likely relates to key differences in how the blastomeres are held together at early cleavage because a more rigorous process was required to allow the separation of blastomeres and even treatment controls did not always develop normally in *P. dumerilii*. Previous authors have suggested that blastomere isolations are not possible in *P. dumerilii* (Fischer and Dorresteijn, 2004) or the related *Alitta virens* (Kostyuchenko and Dondua, 2017) as everything but the trochoblasts cease differentiation. However, Carrillo-Baltodano *et al*. (2025) had success when only removing single blastomeres in *P. dumerilii*, which resulted in partial larvae with defined axes and external features.

### Conclusions

In conclusion, we showed by manipulating blastomeres at early cleavage stages that neural specification is autonomous for the brain and VNC of two annelids, *C. teleta* and *P. dumerilii.* We further showed that neural specification is not coupled to D-V axis organization in the trunk of *C. teleta,* and that episphere and trunk neural tissue is controlled separately. We hypothesize that these processes occur independently in other annelids as well. Uncoupling of neural and D-V axis specification, as well as decoupling episphere and trunk neural specification could have permitted the evolvability of different types of nervous systems and life cycles within annelids, explaining the large diversity of nervous systems in this group.

## Materials and Methods

### Animal care and embryo collection

A colony of *Capitella teleta* Blake, Grassle & Eckelbarger 2009 adults were cultured as previously reported (Grassle and Grassle, 1976; Seaver et al., 2005). Embryos were dissected from the broods of females and cultured in ASW with 60 µg/mL penicillin and 50 µg/mL streptomycin (ASW+PS) (Carrillo-Baltodano and Meyer, 2017; Meyer et al., 2010)

*Platynereis dumerilii* (Audouin and Edwards, 1833) were maintained at the Marine Biological Laboratory, as previously reported (Kuehn et al., 2019). Zygotes were produced by spawning mature worms as described previously. Briefly, mature male and females worms were assembled into a small glass dish until spawning. Zygotes were then rinsed to prevent polyspermy and de-jellied and left to develop until the appropriate stage.

### Blastomere isolations

In all species, embryos of the correct stage were washed three times with ASW+PS. After chemical dissociation (see below for species specifics), the unwanted blastomeres were removed from the rest of the embryo using an eyelash brush in a gelatin- or agarose-coated dish and raised individually in ASW+PS in gelatin- or agarose-coated 24-well dishes until a comparable stage was reached. The delicate embryos were only handled with eyelash brushes and mouth pipettes.

Isolations in *C. teleta* were performed following Carrillo-Baltodano and Meyer (2017). Briefly, embryos at the selected cleavage stage were incubated in 4-well gelatin-coated dishes with Calcium and Magnesium Free Artificial Sea Water (CMF-ASW) (Strathmann, 1987) for 5 min, followed by incubation in 1% protease (Millipore-Sigma cat# P6911) and 1% sodium thioglycolate (Millipore-Sigma cat# T0632) in CMF-ASW for 5 min at room temperature (∼22°C). Embryos were raised individually in ASW+PS for six days at room temperature. ASW+PS was changed daily or not at all in partial and whole-embryo controls, with no changes in survival or health.

In *P. dumerilii*, a stronger dose of thioglycolate was necessary to remove the egg envelope. Embryos were treated as above but on a rocker, and with 1% thioglycolic acid (pH 8, Sigma T6750) instead of sodium thioglycolate. Embryos were raised for 72 h at 18°C in ASW+PS with no water changes to reach a comparable stage.

### Fixation

Animals were relaxed with a 1:1 mixture of ASW plus 0.37 M MgCl_2_ for 5-12 min then fixed with 4% paraformaldehyde (stock 32% PFA ampules from Electron Microscopy Sciences, cat. 15714) in ASW for 30 min for immunolabeling or overnight for whole mount *in situ* hybridization (WMISH). Fixation was stopped with 3 washes in PBS.

### Immunolabeling

After fixation, *C. teleta* embryos were immunolabeled following Meyer *et al*. (2015). Briefly, animals were permeabilized in PBS + 0.1% Triton-X 100 (PBT), blocked in 10% goat serum, and then incubated in primary antibody in block overnight at 4°C. Animals were washed and then incubated with a secondary antibody and 0.1 µg/mL 33342 Hoechst (Millipore-Sigma, cat. B2261) in block overnight at 4°C. Animals were then washed at room temperature with PBT, cleared in 80% glycerol in PBS, and mounted on slides for DIC and fluorescent imaging. Primary antibodies are as follows: 1:600 rabbit anti-serotonin (5HT; Millipore-Sigma, cat #S5545) and 1:800 mouse anti-acetylated tubulin (clone 6-11B-1, Millipore-Sigma, cat #T6793). Secondary antibodies were as follows: 1:300 goat anti-mouse F(ab’)_2_ FITC (Millipore-Sigma, cat #F8521) and 1:600 sheep anti-rabbit F(ab’)_2_ Cy3 (Millipore-Sigma, cat #C2306). 1:100 BODIPY FL-Phallacidin (Life Technologies, cat #B607; stock concentration 200 Units/mL in methanol) was included with secondary antibodies.

In *P. dumerilii,* immunolabelling followed the above, except an additional proteinase K digestion step was added for permeabilization (2 min in 100 μg/mL proteinase K), followed by a 20 min post fixation in 4% PFA.

### Whole Mount *In Situ* Hybridization (WMISH)

WMISH was conducted as described previously (Seaver et al., 2001). After fixation, animals were serially dehydrated in methanol and stored at -20 °C. Animals were hybridized for a minimum of 72 h at 65 °C with 1 ng/μl of each probe. Spatiotemporal RNA localization was observed using an NBT/BCIP color reaction. The color reaction was stopped using 3 washes of PBS + 0.1% Tween-20. After WMISH, animals were labeled with 0.1 μg/ml Hoechst 33342 (Sigma-Aldrich, cat. B2261), cleared in 80% glycerol in PBS, and mounted on slides for DIC and fluorescent imaging.

RNA probes as used in (Boyle and Seaver, 2008; Meyer and Seaver, 2009), and with 2 ng/µL of DIG-labeled anti-sense *Ct-bmp5-8* RNA probe used in Corbet (2016).

### Imaging

Immunolabelled and WMISH animals were imaged using DIC optics on an AxioImager M2 compound microscope (Zeiss) coupled with an 18.0 megapixel EOS Rebel T2i digital camera (Canon) or with an AxioCam MRm rev.3 camera (Zeiss) and Zen Blue software (Zeiss). DIC images were rendered in Helicon Focus (HelSoft). Z-projections were obtained from the ZenBlue stacks using Fiji (Image J2, NIH). Contrast and brightness of immunolabelling and WMISH images were edited in Photoshop CC, and figure panels were constructed using Illustrator CC (Adobe Systems, Inc.).

## Acknowledgements

The authors would like to thank the staff at the Marine Biological Laboratory for their strong support of early researchers.

## Author contributions

Following the CRediT taxonomy, ACB, NPM, and NBW conceived of the study, ACB, NPM, NBW, JDS, SD collected data, ACB and NBW prepared visualization, ACB and NBW wrote the manuscript, ACB, NPM, NBW, JDS, and BDO edited the manuscript, and NM and BDO provided resources.

## Data and resource availability

The datasets used and/or analyzed during the current study are available from the corresponding author upon request.

